# Single organoid RNA-sequencing reveals high organoid-to-organoid variability

**DOI:** 10.1101/2021.11.22.469588

**Authors:** Kristin Gehling, Swati Parekh, Farina Schneider, Marcel Kirchner, Vangelis Kondylis, Chrysa Nikopoulou, Peter Tessarz

## Abstract

Over the last decades, organoids have been established from the majority of tissue resident stem and iPS cells. They hold great promise for our understanding of mammalian organ development, but also for the study of disease or even personalized medicine. In recent years, several reports hinted at intraculture organoid variability, but a systematic analysis of such a heterogeneity has not been performed before. Here, we used RNA-seq of individual organoids to address this question. Importantly, we find that batch-to-batch variation is very low, even when prepared by different researchers. On the other hand, there is organoid-to-organoid variability within a culture. Using differential gene expression, we did not identify specific pathways that drive this variability, pointing towards possible effects of the microenvironment within the culture condition. Taken together, our study provides a framework for organoid researchers to properly consider experimental design.

## INTRODUCTION

Organoid cultures have been present in modern research laboratories for over a decade and are thought to bridge the gap between 2D and 3D-tissue culture (Broutier et al. 2017; Aizarani et al. 2019; Lancaster and Huch 2019). Organoids can be derived either from pluripotent cells, such as embryonic or induced pluripotent stem cells, but also from tissue-resident stem and progenitor cells (Prior et al. 2019). In particular, iPSC-derived organoids can give rise to remarkably complex structures (Takebe et al. 2013). Recently, gene regulatory network analysis using CellNet (Cahan et al. 2014) in combination with CRISPR-Cas based engineering was used to generate complex organoids (Velazquez et al. 2021). While high complexity as well as disease modelling can be nowadays derived in iPSC-derived organoids, they lack epigenetic information of the tissue of origin, which might hamper analysis of complex states, such as cancer. Thus, besides complex multilineage organoids, 3D structures have been derived from tissue-resident progenitors (Broutier et al. 2017).

In the case of the liver, organoids are based on hepatocytes and cholangiocytes. Hepatocyte-derived organoids, so called ‘Hep-Orgs’ consist mostly of progenitors and hepatocytes (Hu et al. 2018). In contrast, ‘Chol-Orgs’ are derived from EPCAM+ or Lgr5+ biliary epithelial cells and have the potential to differentiate into either hepatocytes or cholangiocytes (Huch et al. 2013). Upon *in vitro* differentiation, cholangiocyte-derived organoids will give rise to functional cells that display hepatocyte characteristics, like increased glycogen storage, LDL uptake or albumin secretion. These hepatocytes can be transplanted into liver-damaged Fah-/- mice where they contribute to liver tissue and thus, prolonged life-span (Huch et al. 2013). This murine model system is frequently used in liver biology and allows repopulation with hepatocytes.

In recent years, one focus was the establishment of 3D cultures from various organs and nowadays organoids can be grown representing virtually any tissue. The vision in ongoing consortia is to exploit the tissue-like features of organoids to understand the development of human disease (Rajewsky et al. 2020) and thus, similar to an organismal atlas, organoids are also currently profiled to generate an overview of cell types present as part of the human cell atlas project (Bock et al. 2021).

To date, organoids represent the model system, which most closely resembles the tissue of origin. The multicellular nature of organoids makes them a sophisticated but variable model, which displays heterogeneity (Lancaster and Knoblich 2014) and can strongly depend on many extrinsic factors, such as culture conditions (Criss et al. 2021). However, we still need to better understand the drivers of batch-to-batch and organoid-to-organoid variability within the same culture. To address these questions, we profiled single organoids from 4 different batches and passage numbers (Figure 1A). To allow an easier isolation of single organoids, we initially set up shaking cultures from organoids derived from adult livers and compared their gene expression programs with those of the same organoids growing in domes. Gene expression profiling revealed a striking change in the transcriptional program towards a more progenitor-like state, with an increase in proliferative terms. Next, we isolated single, intact organoids, which were macroscopically evaluated before RNA-extraction and library preparation. This approach resulted in the generation of 35 single organoid libraries from four organoid batches. The batch-to-batch variation was very low - even between batches generated by different scientists, with passage number being the most likely reason for gene expression differences. However, the variability between organoids within a given batch was much larger, but was not determined by size or overall cell cycle state. Taken together, we provide a resource that addresses confounding factors in organoid culture, which will hopefully help the community with their experimental design.

**Figure 1:**
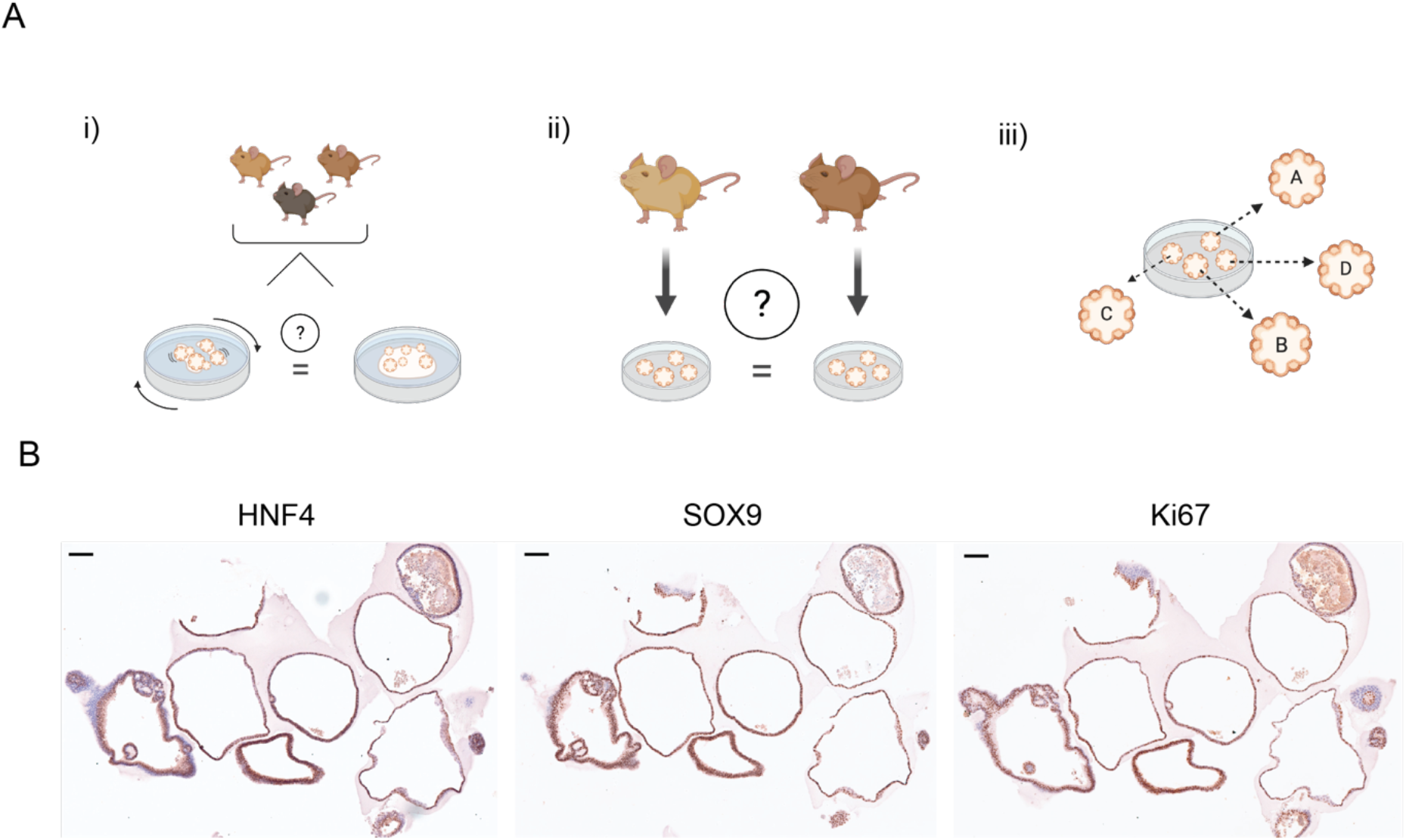
Rational and Setup of the Study. A) This Study aims to investigate organoids i) grown in a shaking culture and evaluate them as an alternative to dome culture, ii) generated from different animals to interrogate batch effects and iii) analysis of single organoids to assess the heterogeneity within a culture. B) Immunohistochemistry of the same sample of young organoids for HNF4, SOX9 and Ki67. Scale bar is 100μm.

## MATERIALS AND METHODS

### Initiation of liver duct-derived organoid cultures

Three-month old C57BL/6N mice were maintained in the mouse facility of Max Planck Institute for Biology of Ageing and sacrificed according to approved ethical guidelines (granted by the Landesamt für Natur, Umwelt und Verbraucherschutz Nordrhein-Westfalen).

Cultures for comparing the culture methods were made by pooling digested material from different animals each (Pool A = 2 mice; Pool B = 4 mice; Pool C = 3 mice). Organoid cultures for the heterogeneity experiment (Set 1 - Set 4) were established from the liver of only one mouse for each set.

Livers were excised postmortem and digested according to the manufacturer’s protocol (HepatiCult, StemCell Technology, 06030) with a few modifications. A total of three digestion cycles were needed to dissolve the 3-5 mm pieces of liver tissue. For duct isolation, the 70 μM strainer was omitted, and the pooled supernatant only passed through a 35 μM cell strainer. The strainer was reversed onto a pre-wetted falcon tube and 10 ml cold Advanced DMEM/F-12 added to release the hepatic ducts from the strainer. To ensure the detachment of big fragments, the bottom of the filter was scraped with a P1000 pipette and the remaining fragments were repeatedly collected with a total of 2 ml Advanced DMEM/F-12.

#### Culture in Matrigel domes

The pelleted ducts were cultured in 30 μl Matrigel domes as described in the Supplementary Protocol for Mouse Hepatic Progenitor Organoid Culture (Catalog #06030). Organoid structures arose within 6 days. Organoids were maintained in a 37 °C incubator at 5% CO2 and 20% O2. The medium was changed every 2 days and cultures passaged every 5-7 days with mechanical dissociation of the Matrigel.

#### Suspension cultures in 10% Matrigel

For initiating organoid cultured in a dilute suspension culture, 50 μl of a 1:10 Matrigel/complete HepatiCult mixture was mixed with the duct pellet and pipetted into one well of a cooled 12-well-plate, already containing 950 μl of the Matrigel/HepatiCult mixture. The cultures were maintained on an orbital shaker at 80 r.p.m. in a 37 °C incubator, 5% CO2, 20% O2. Every 3-4 days, the organoids were passaged by mechanical breakdown of the Matrigel.

### Selection of individual organoids

Organoids were maintained as dilute suspension cultures. Set 1, 2, and 3 were seeded onto 24-well plates at 5,4, and 3 days before extraction. Set 4 was seeded onto a 12-well plate 3 days before mRNA extraction. A sterile and RNAse-free work environment was organized in the best manner. The tip of P200 filter tips was cut with a sterile razor to allow the pipetting of organoids in a volume of 10-20 μl without disturbing the lumen. Using a Leica M80 Stereo Microscope, individual organoids were carefully transferred into a neighboring well with DPBS and subsequently added to a cooled 24-well plate with 10 μl droplets of DMEM/F-12, resulting in one organoid per well.

### Immunohistochemistry

Organoids were fixed *in situ* in 4% PFA for 1 hr at RT, washed twice with 1XPBS and isolated by mechanical disruption of the matrigel. The organoids were then processed for paraffin embedding. Sections of paraffin-embedded samples were deparaffinised by immersion of the slides into the following buffers; 20 min in Xylol, 2 min with 96% EtOH, 2 min with 75% EtOH, 2 min with 1X PBS and washed three times with H_2_O for 5 min each. Endogenous peroxidase was quenched by immersion for 15 min in peroxidase blocking buffer (0.04 M NaCitrate pH 6.0, 0.121 M Na2HPO4, 0.03 M NaN3, 3% H_2_O_2_). After three washes with tap water, slides were subjected to heat-induced epitope retrieval with 10 mM NaCitrate, 0.05% Tween-20, pH 6.0, washed 5 min with 1X PBS, blocked 60 min with Blocking buffer + 160 μl/ml AvidinD and incubated with primary antibodies diluted (1:400 Ki67, 1:200 SOX9, and HNF4a) in blocking buffer + 160 μl/ml Biotin overnight at 4°C. After three 5 min washes with PBST the samples were incubated with the secondary antibody 1:1000 diluted in blocking buffer for 1 h at room temperature, followed by three 5 min washes with PBST and incubation for 3 0min with 1x PBS + 1:60 Avidin D + 1:60 Biotin. After three 5 min washes with PBST the samples were stained with 1 drop of DAB chromogen in 1 ml Substrate buffer, washed with 1X PBS and counterstained with Hematoxylin for 4 min, washed with tap water and dehydrated 1min in each buffer; 75% EtOH, 96% EtOH, 100% EtOH, Xylol and mounted with Entellan.

### Microscopy

Immunohistochemistry images were taken using the slidescanner Hamamatsu S360 and analysed with the NDP.view2 software. Images were taken with the EVOS FL Auto 2 Imaging System in standard brightfield with a 4x/0,13 NA or a 10x/0,25 NA objective. To calculate the organoid area, acquired raw files were analyzed with an automated macro in FIJI using the following steps: Gaussian Blur with radius of sigma =3, Auto Threshold method = MaxEntropy followed by the “Fill Holes” function of the binary mask. Subsequent “Analyze Particles” delivered the desired areas.

### mRNA extraction

The mRNA was extracted with the Dynabeads mRNA DIRECT Kit by Fisher Scientific (#61011) with a few modifications and self-made buffers. For each sample (i.e., single organoid) 10 μl of resuspended beads were transferred to a 2 ml low-binding tube and placed on a DynaMag-2 magnet stand. The supernatant was discarded, the magnet removed and beads were resuspended in 50 μl room temperature Lysis/Binding Buffer. With a volume of 150 μl room temperature Lysis/Binding Buffer, each organoid was transferred to a 1.5 ml low-binding tube already containing the equivalent amount of buffer. The content was pipetted up and down 5 times to allow lysis. The following steps were performed according to the manufacturer’s instructions. The mRNAs were normalized for library preparation input by measuring actin (Actb Fw: 5’-CAGCTTCTTTGCAGCTCCTT Rv: 5’-CACGATGGAGGGGAATACAG) expression via quantitative PCR. The Luna Universal Probe One-Step RTqPCRKit by New England Biolabs (#E3005S) was used to combine reverse transcription (RT) with quantitative polymerase chain reaction (qPCR). A scaling factor for the mRNA for consecutive library preparation (protocol below) was calculated with the following formula: c_qmax_-c_qmean_ =2 ^scaling factor^. Cqmean was calculated from two independent qPCR runs. Cqmax was set to 21. The maximum input volume for reverse transcription is 6.4 μl, thus 6.4 was divided by each sample’s scaling factor and yielded a normalized amount of input mRNA.

### RNA-seq

RNA libraries were created as previously described (Allmeroth et al. 2021). In brief, equal amounts of mRNA per sample were used for cDNA synthesis with Maxima H Minus reverse transcriptase (Thermo Fisher Scientific). During reverse transcription, unique barcodes including unique molecular identifiers (UMI) were attached to each sample. After cDNA synthesis, all samples were pooled and processed in one single tube. DNA was purified using AmpureXP beads (Beckman Coulter) and the eluted cDNA was subjected to Exonuclease I treatment (New England Biolabs). cDNA was PCR-amplified for 12 cycles and subsequently purified. After purification, cDNA was tagmented in 10 technical replicates of 1 ng cDNA each using the Nextera XT Kit (Illumina), according to the manufacturer’s instructions. The final library was purified and concentration and size were validated by Qubit and High Sensitivity TapeStation D1000 analyses. Paired-end sequencing was performed on Illumina NovaSeq 6000. Fastq files were processed with zUMIs (version 2.9.5) using its miniconda environment (Parekh et al. 2018) with STAR index 2.7 (Dobin et al. 2013), samtools (version 1.9) (Li et al. 2009) and “featureCounts” from Rsubread (version 1.32.4) (Liao et al. 2013). The reads were mapped to *Mus musculus* (mm10) with Ensembl annotation version GRCm38.93. Libraries were down-sampled within zUMIs, depending on library size variability. Downstream computational analysis was conducted in R (version 3.6.3). The count matrix was normalized and filtered with edgeR (version 3.28.1) (Robinson et al. 2010) using “filterByExpr” with the min.count = 3. For differential gene expression analysis, the limma-voom approached by limma (version 3.42.2) (Ritchie et al. 2015) was used with a pipeline including linear model fit (lmFit) and p-value adjustment for multiple testing (“topTableF” with adjust.method = ‘BH’, “decideTests” with method = ‘global’). Obtained sets of genes were further analyzed, e.g. through gene enrichment analysis with MetaScape (Zhou et al. 2019). Intersections were visualized with UpsetR (version 1.4.0) (Conway et al. 2017), heatmaps created with pheatmap, version 1.0.12 (Kolde 2019) and cell cycle analysis with cyclone (Scialdone et al. 2015). Results were plotted with ggplot2, version 3.3.3 (Wickham 2011).

## RESULTS

### Heterogeneity within organoid culture

To address overall heterogeneity within organoid dome cultures, we initially performed several stainings for progenitor or proliferation markers. While the majority of cells within organoids were positive for Sox9, there was a variable amount of Ki67 and HNF4 positive cells from one organoid to another (Figure 1B). In the context of heterogeneity, it is also important to highlight that even within a single organoid, there can be a high degree of variability as seen in the individual stainings. These results prompted us to investigate the heterogeneity between organoids further and to understand if there are specific pathways that might explain the observed variability. We have additionally tested the reproducibility of organoid generation.

### Comparison of dome vs shaking culture

To enable simple isolation of individual organoids, we wanted to set up culture conditions, in which cells would grow in a low percentage of Matrigel that would be amenable to pipetting individual organoids. To compare the low percentage Matrigel cultures with the classic cultures of organoids in domes, we set up an experimental design to marginalize effects of individual mice, yet evaluate the reproducibility with different biological replicates. Therefore, organoids were generated from pooled murine liver tissue of different animals. In total, ductal organoids were initiated from three replicates of different three-month-old mice (Pool 1: two animals; Pool 2: four animals; Pool 3: three animals). After establishment of organoid cultures, organoids were then split to continue growth in Matrigel domes, or were seeded at 10% Matrigel concentration using a shaking platform (see Methods for details). We assessed the gene expression program for both conditions using 3’-end RNA-seq, which was subsequently analysed using ZUMIs (Parekh et al. 2018).

After filtering for an adjusted p-value below 0.05, 3225 genes were found differentially expressed between dome and suspension cultures, of which 1635 were down and 1590 upregulated (Figure 2A, Supplementary Table 1). The heatmap of differentially expressed genes showed a clear separation between the two conditions and suggested higher variability between samples in the dome cultures (Figure 2B). To investigate possible pathways and functions behind differentially expressed genes, we performed gene ontology enrichment using Metascape on genes with a fold-change >1, which resulted in 1015 up- and 1172 down-regulated genes (Zhou et al. 2019). Most of the terms enriched in dome-cultured organoids were associated with metabolic terms (genes included *Cyp2b10, Cyp2c29, Cyp2j6*; Figure 2C), while suspension cultures showed enrichment in terms that were connected to proliferation, such as ‘mitotic cell cycle process’ or ‘DNA replication’ (*Cdk1, Cdk11b, Cdc45, Cdk4*; Figure 2D, Supplementary Table 1). Taken together, the observed gene expression differences suggested that organoids grown within a Matrigel dome represented a more mature, liver-like state than those within the shaking cultures, which was dominated by proliferative terms.

**Fig. 2:**
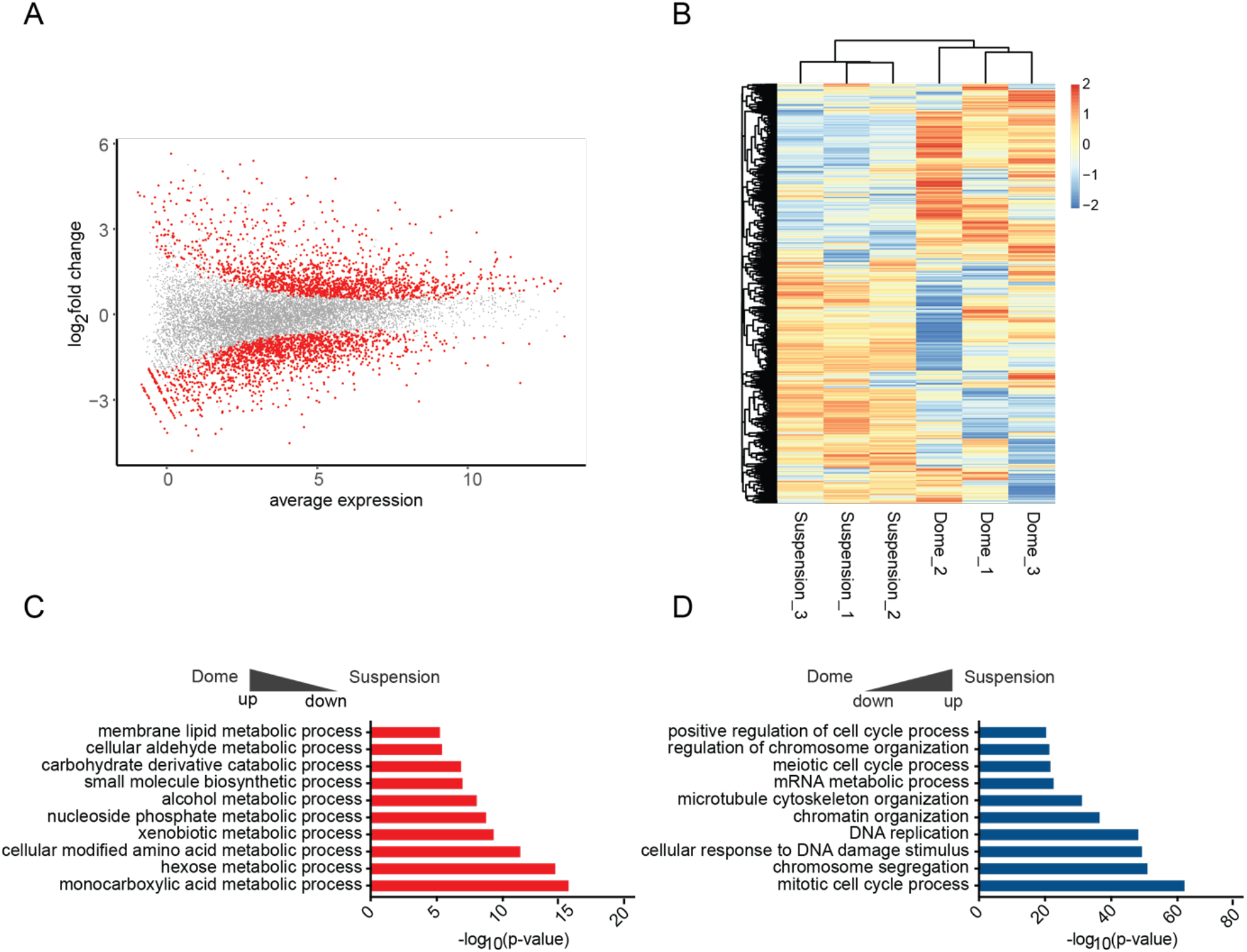
Differential gene expression analysis of dome and suspension cultured ductal-derived organoids. A) MA-plot showing the log ratio and average expression values of DEGs. Significant genes after p-value adjustment are displayed in red. B) The heatmap shows the normalized expression values generated by limma-voom for the fraction of significant DEGs. Heatmap data is clustered by both row and column, and scaled by row. C), D) Metascape enrichment analysis of DEGs comparing dome and suspension culture. The top 20 bar graphs are shown, and respective clusters are coloured by p-value enrichment.

### Passage Number is the strongest contributor to gene expression changes in organoid cultures

To assess the heterogeneity of organoids, new cultures were established from four three-month old male mice. Each culture originated from the tissue of one animal. After establishment of organoid culture in domes, cells were subsequently seeded in suspension culture for at least two passages. Individual organoids from different wells of the same biological replicate were carefully transferred with a pipette into a new 24-well plate. Only non-apoptotic organoids without a dark necrotic lumen, were selected, including a variety of sizes.

Before mRNA extraction, images were taken and organoids were further selected based on singularity per well and undamaged lumen. Representative organoids are displayed in Figure 3A. The images were used to calculate the 2D surface area (Supplementary Figure 1A, B). Organoids from Set 1 were analysed after passage 11 and displayed a similar morphology within the set regarding the evenness of the lumen compared to all other sets. Sets 2, 3 were harvested at passage three and Set 4 at passage four. In total, we generated 42 RNA-seq libraries across the four sets, of which 35 libraries passed quality control and filtering for a minimum library size of four million reads. In order to visualize the variance within the dataset, principal component analysis (PCA) was performed (Figure 3B,C). The library size of each sample is annotated in counts per million and represented by the size of the dot. Importantly, library size differences (though present) did not drive separation between the individual organoids. Interestingly, while organoids within sets 2-4 clustered closely together, set 1 formed a distinct cluster (Figure 3B,C). As the major difference between set 1 and sets 2-4 was a higher passage number, this indicated that changes occurred in these organoids upon prolonged culturing. To investigate this potential passage effect further, we performed differential expression testing between all sets. The analysis was performed using the linear models-approach within limma (Law et al. 2014; Ritchie et al. 2015). As this study did not involve a simple control-treatment design, the 4 different sets were juxtaposed within six comparisons. The expression values of significantly expressed genes (adj. p-value < 0.05) were subsequently displayed in a heatmap (Figure 3D, Supplementary Table 2) and data were clustered by sample (i.e. column). The heatmap replicated the trend seen in the dimensionality reduction analysis above and showed most of the set 1 to be clustering in a separate branch. While the other three sets appear ordered according to their passage number, suggesting that indeed, gene expression was dominated here by passage number. To address the changes in gene expression on a more functional level, we first compared the number of differentially expressed genes across the sets (Supplementary Figure 2). Not surprisingly, comparisons with set 1 showed the strongest differential expression. We then chose the intersection between these comparisons with set 1 and performed GO term enrichment analysis for upregulated genes in set 1 (Figure 3E). The enrichment map of GO terms pointed towards a strong enrichment for liver-specific functions, indicating that prolonged culture leads to a more differentiated phenotype. The enrichment for a more progenitor-like state is evident in the GO analysis of down-regulated genes (Figure 3F), in which proliferative and developmental terms were dominating.

**Fig. 3:**
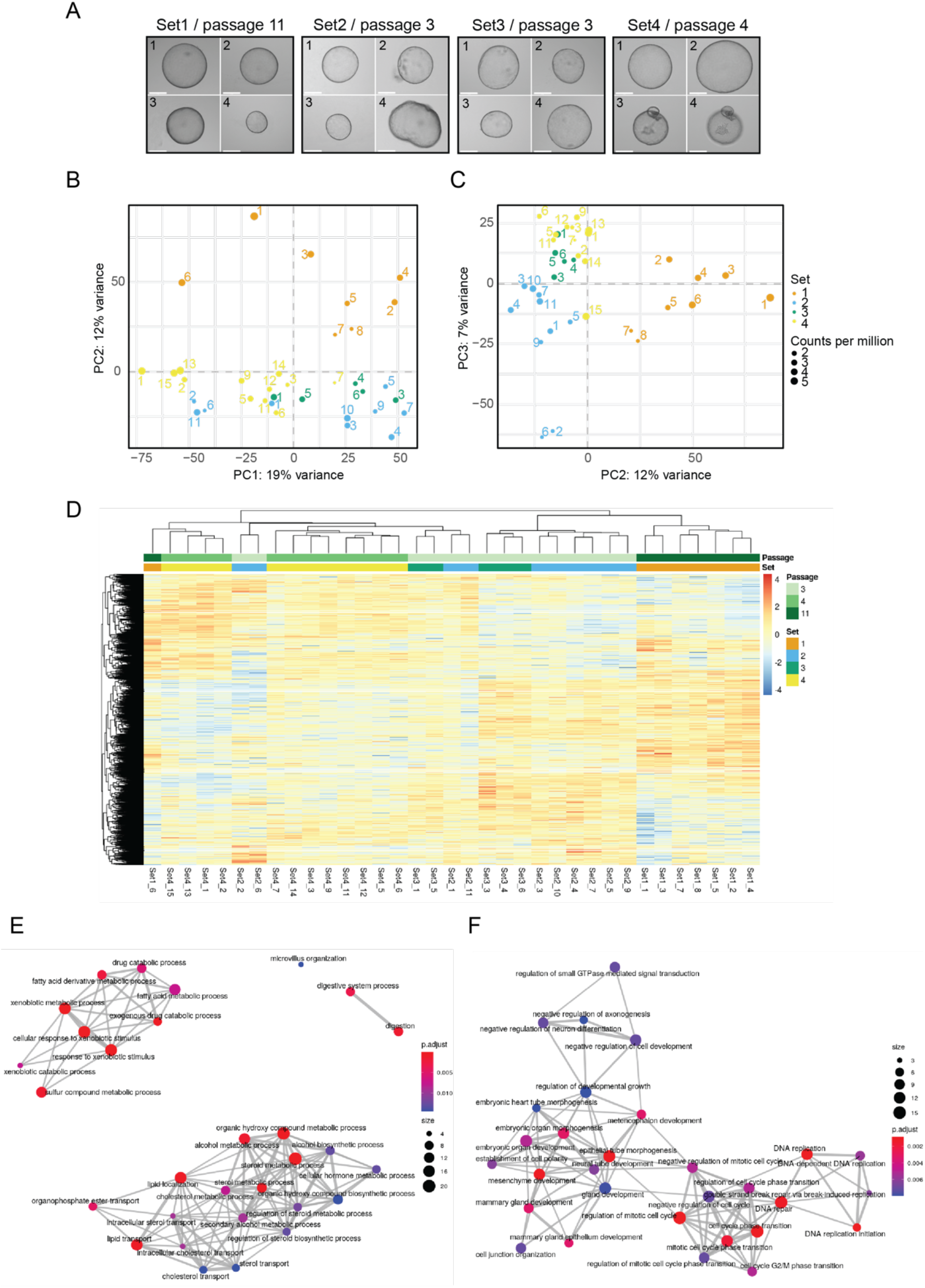
Analysis of single organoids. A) Representative microscopy images taken from individual organoids. The passage number at the day of sorting is indicated for each set. Sample numbers are written in the top left corner of each image, and the scale bar represents 275 µm. B), C) The grouping of organoids after dimensionality reduction by principal component analysis (PCA) for the first two PCs and PC 2 and 3. The distribution of variance among the PCs is plotted along, with each proportional contribution to the variance in the axis labels. The dot size indicates library size in counts per million. D) Heatmap of expression values after DEA. Significant genes after p-value adjustment by BH procedure are clustered by both row and column, and scaled by row. E) Enrichment map showing GO term analysis of upregulated DEGs shared against set 1. F) The intersection of downregulated DEGs from Set2vs1, Set3vs1 and Set4vs1 plotted as enrichment map.

### Organoid-to-organoid variability

We assessed the heterogeneity within each set using PCA (Figure 4A-D, which indicated a high degree of variability between individual organoids. To understand in more detail what drove the separation, we analysed the genes driving the separation of PC1 and PC2 for each set (Supplementary Table 3). However, we did not identify any enriched term for the differentially expressed genes, indicating that there were no changes in specific pathways between the individual organoids. As the unbiased approach did not yield any specific pathway, we took a candidate-based approach. We chose marker genes that report proliferation status, response to Wnt signalling or differentiated vs. progenitor-like state (Figure 4E). The heatmap confirms the high level of heterogeneity between the individual organoids. It also corroborates the finding that organoids from set 1 have a more hepatic-like state compared to the others. Interestingly, even within a given culture like set 2, organoids can express high levels of mature hepatic markers (e.g. *Fah* and *Ldlr*), while others were enriched in ductal and progenitor markers (e.g. *Notch2, Lgr5* or *EPCAM*), indicating that organoids within one culture can lean towards two different fates.

**Fig. 4:**
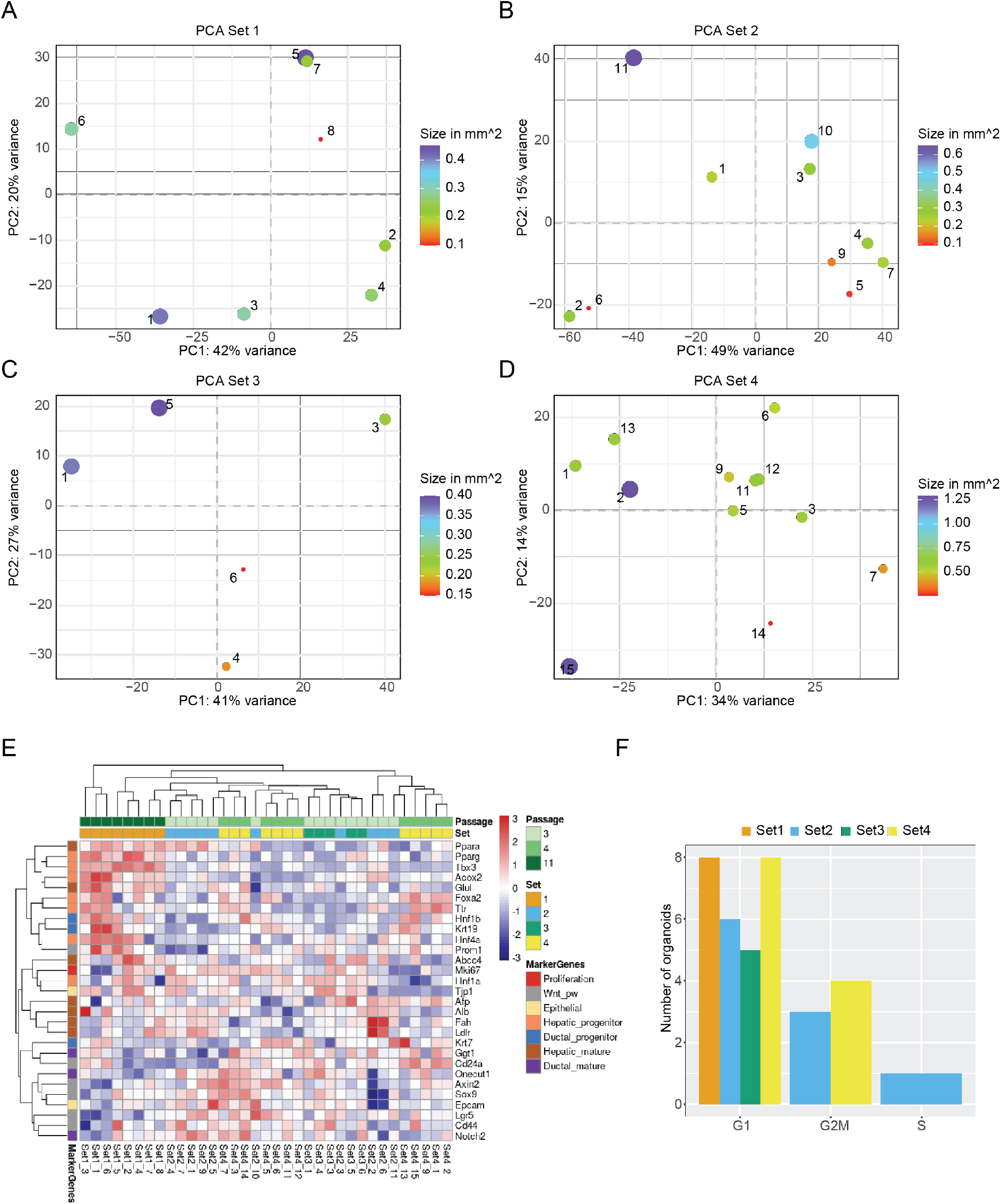
Inter-organoid heterogeneity. A) - D) Projection of organoids within each set after dimensionality reduction by PCA, including the annotation of each organoid’s size as dot size. The variance for each PC is indicated in the axis labels. E) Heatmap showing the expression values for marker genes after limma-voom normalization, with a minimum count = 1. Rows are annotated by official gene symbols and colours indicating marker for proliferation (red), Wnt pathway (grey), epithelial (yellow) as well as progenitor and mature cell types for hepatic (orange, brown) and ductal (blue, violet). Data is clustered hierarchically by row and column and scaled by row. F) Bar graph showing number of organoids for each cell cycle phase, coloured by set, after analysis with cyclone.

To understand whether the size of an organoid might impact gene expression programs, we went back to the PCA analysis, in which the size in which the size of the organoid is indicated (Figure 4A-D). However, no clear relationship between transcriptome and size was observable. To confirm this result, we plotted the loadings of the first two principal components against the size of the organoids and fitted a regression line to the data (Supplementary Figure 3A,B). While in some instances there was a good correlation between size and the PC loadings, this was not the case in general. In conclusion, organoid size does not seem to be a driver of the individual transcription programs within single organoids. Finally, we evaluated whether the overall proliferative state of organoids might contribute to the differences in transcriptional states. As a proxy for the level of proliferation, we performed a cell cycle analysis using Cyclone (Scialdone et al. 2015). Cyclone is a machine learning based approach allowing cell cycle stage prediction based on a reference transcriptome. Here, a sample is assigned to G1 or G2M, when it reaches the threshold of 0.5 for the particular phase. If both G1 and G2M scores stay below 0.5, the sample will be categorized as S-phase. The majority of organoids were assigned to G1 phase (Figure 4F). While some organoids were predicted to fall more into G2M phase, this assignment did not correlate with the clustering in the PCA. Still, most organoids showed variability in the cell cycle score. In conclusion, proliferative states of individual organoids will contribute to the organoid-to-organoid variability as well as size, but these two parameters alone were not able to predict the stark differences in gene expression programs.

## DISCUSSION AND CONCLUSION

Organoid-to-organoid variability has been observed and reported before for epithelial organoids (Hof et al. 2021) and is particularly prevalent in organoids recapitulating high tissue complexity, such as brain organoids (Quadrato et al. 2017; Velasco et al. 2019). Recent studies have analysed in depth the effect of different culture conditions and treatments in gene expression variability (Criss et al. 2021) and the donor batch effect of different donors on cultures of human gut organoids (Mohammadi et al. 2021). Here, we report that also less complex organoids, derived from genetically identical mice show a high degree of variability from organoid to organoid. Nevertheless, reproducibility (as measured in bulk assays) was high between several batches of organoids.

While organoid-to-organoid variability was obvious and marker gene analysis suggested a variety of different cellular states between the organoids, the underlying reason for the variability was less clear. Organoid size and proliferation state - as delineated from predicted cell cycle stage - contributed to the overall variability, but their impacts were not large enough to fully explain the extent of variability. Given the contribution of culture conditions on variability as seen in the shaking organoid culture, it is reasonable to assume that intra-well conditions might be a strong driver of culture variability (Snijder and Pelkmans 2011). During culturing, assemblies of organoids of various sizes and numbers can be observed, as well as single organoids. Thus, cell-to-cell or cell-to-Matrigel contacts will be different in each scenario and might change the underlying transcriptional program. This might ultimately lead to different signalling events as well. Taken together, in order to grasp biological meaningful signals, scientists need to include multiple technical replicates from organoid cultures of different biological hosts. In addition, depending on the question, specific culture conditions, passaging and culturing time is an important consideration as it can change the cellular state within the organoids.

## Acknowledgements and Funding Information

We would like to thank members of the Tessarz lab and Achim Tresch for discussions. We are particularly grateful to Niklas Kleinenkuhnen for critical reading of the manuscript. We would like to thank Pauline Camelot for help with setting up some of the organoid cultures. We are indebted to the FACS and Imaging facility of the MPI for Biology of Ageing and the Imaging Facility of CECAD, University of Cologne for help with imaging. Sequencing was performed at the Sequencing Core Facility of the MPI for Molecular Genetics in Berlin, Germany. This work was funded by the Max Planck Society (to P.T.), the BOOST program of the Max Planck Society (to C.N.) and a Cologne Graduate School for Ageing Research Master fellowship (to K.G.).

## AUTHOR CONTRIBUTION

Conceptualisation: K.G., C.N., P.T.; Methodology: K.G., M.K., C.N.; Investigation: K.G., F.S., V.K., C.N.; Formal Analysis: K.G., S.P.; Supervision: C.N., P.T.; Funding Acquisition: P.T.; Project Administration: C.N., P.T.; Writing of Manuscript: C.N., P.T., with input from all authors

## DATA AVAILABILITY

All RNA-seq data presented here is available at Gene Expression Omnibus, accession number:

## CONFLICT OF INTEREST

The authors do not declare any conflict of interest.

## SUPPLEMENTARY FIGURES

**Supplementary Figure 1:**
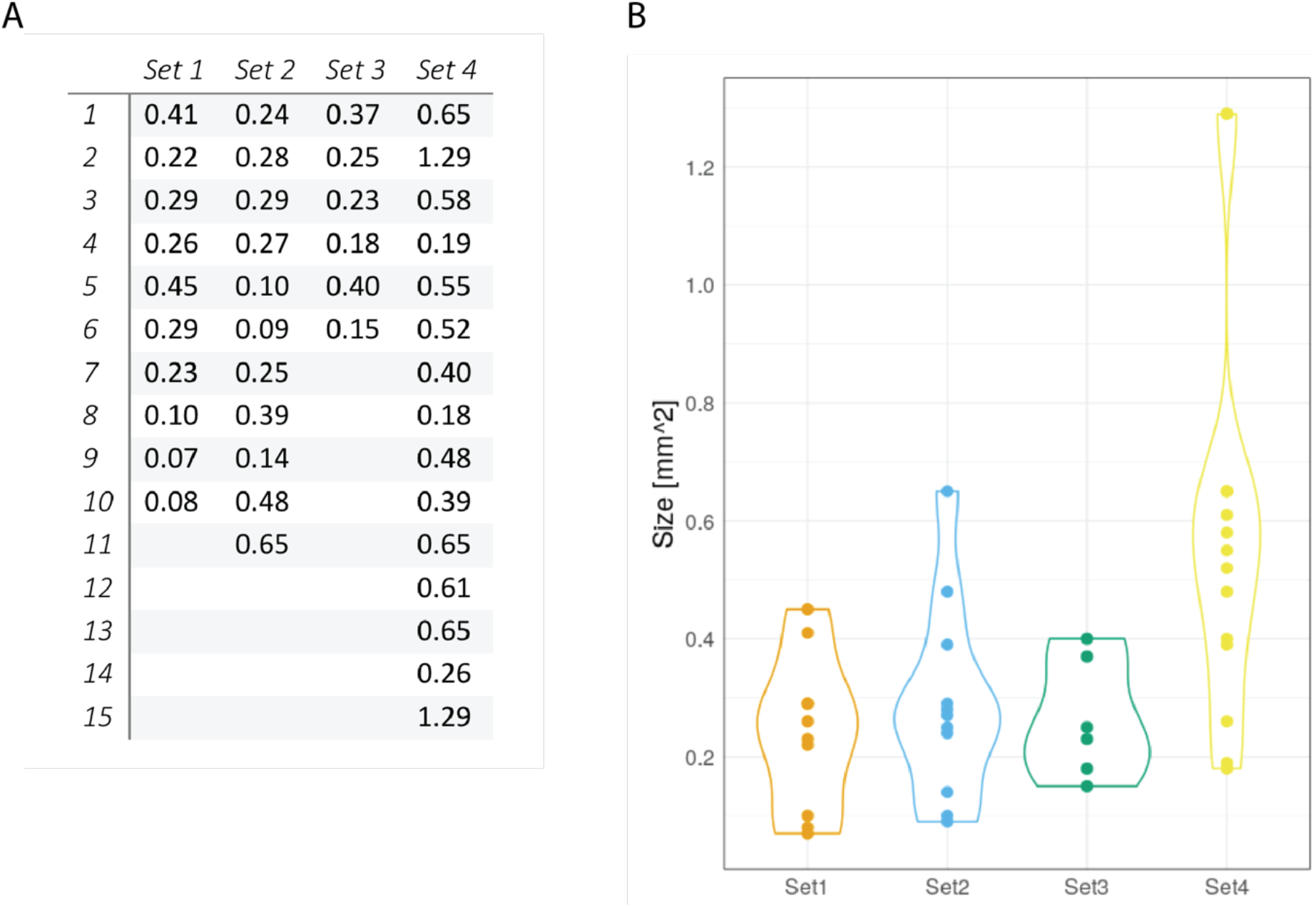
Sizes of individual organoids. Sizes of individual organoids in square mm measured from 2D images with FIJI as table (A) and graphic (B).

**Supplementary Figure 2:**
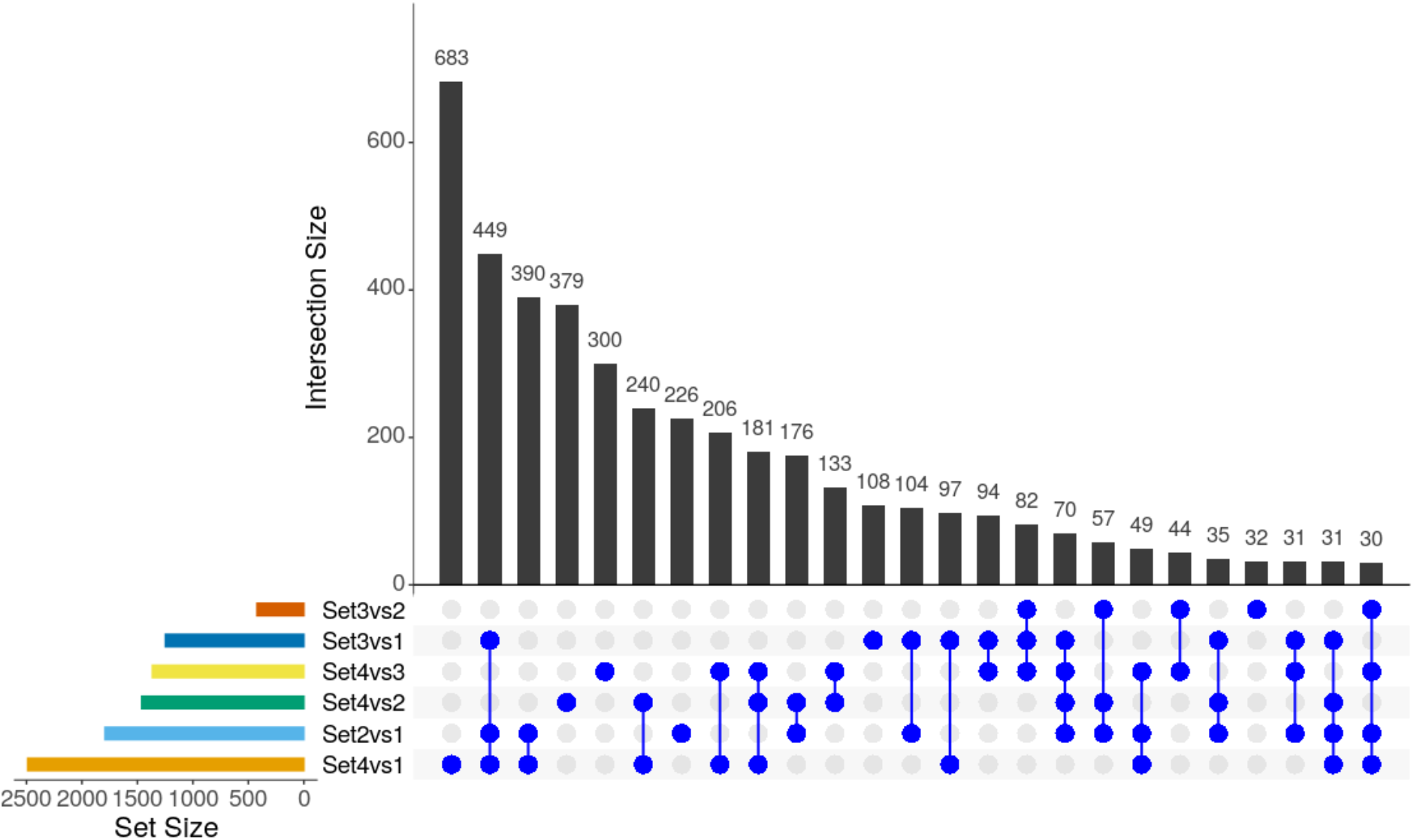
Differential expression analysis by limma-voom approach across sets. A) The upset plot displays the overlap of differentially expressed genes (DEGs) between comparisons. The coloured horizontal bar graphs indicate the number of DEGs for each comparison, respectively. The black vertical bar graphs visualize the intersection size of DEGs and the blue dots represent contributing comparisons from the DE analysis. Only the first 25 intersections are plotted.

**Supplementary Figure 3:**
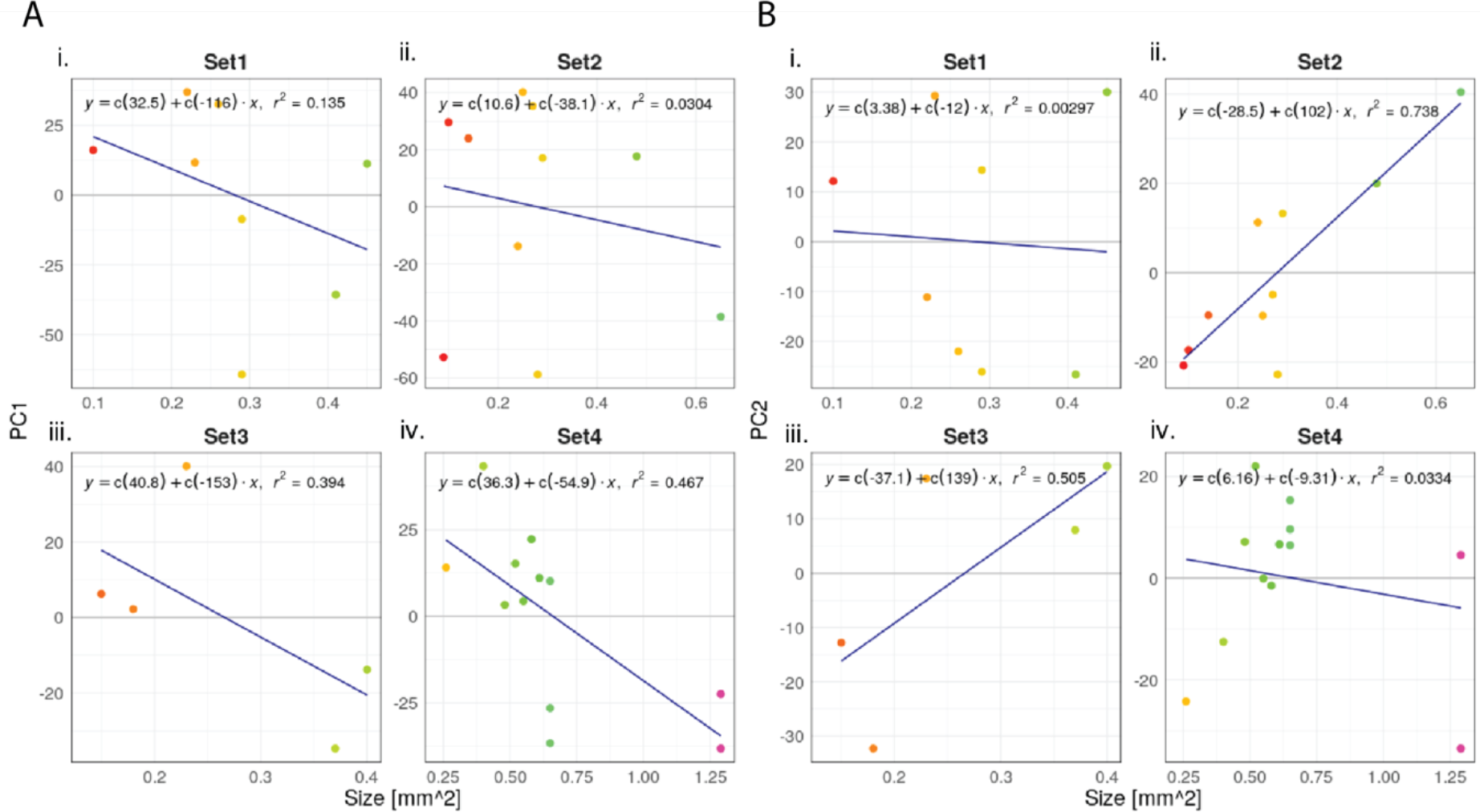
Investigation of a potential correlation between organoid size and principal components. Principal component (PC) 1 in (A) and PC 2 in (B) are plotted for Set 1 – Set 4 in i) -iv), respectively. Organoids are coloured according to their similarity in size. A regression line was fitted including respective formulas and r-squared values. F-statistics for PC1: Set 1 = 0.03711, Set 2 = 0.6302, Set 3 = 0.2571, Set 4 = 0.01428; PC2: Set 1 = 0.898, Set 2 = 0.001454, Set 3 = 0.1787, Set 4 = 0.5695.

## Notes

### Competing Interest Statement

The authors have declared no competing interest.

